# SMGen: A generator of synthetic models of biochemical reaction networks

**DOI:** 10.1101/2021.07.29.454343

**Authors:** Simone G. Riva, Paolo Cazzaniga, Marco S. Nobile, Simone Spolaor, Leonardo Rundo, Daniela Besozzi, Andrea Tangherloni

**Author notes:** Correspondence (S.G.R.); (A.T.).

## Abstract

Several software tools for the simulation and analysis of biochemical reaction networks have been developed in the last decades; however, assessing and comparing their computational performance in executing the typical tasks of Computational Systems Biology can be limited by the lack of a standardized benchmarking approach. To overcome these limitations, we propose here a novel tool, named SMGen, designed to automatically generate synthetic models of reaction networks that, by construction, are characterized by both features (e.g., system connectivity, reaction discreteness) and non trivial emergent dynamics of real biochemical networks. The generation of synthetic models in SMGen is based on the definition of an undirected graph consisting of a single connected component, which generally results in a computationally demanding task. To avoid any burden in the execution time, SMGen exploits a Main-Worker paradigm to speed up the overall process. SMGen is also provided with a user-friendly Graphical User Interface that allows the user to easily set up all the parameters required to generate a set of synthetic models with any user-defined number of reactions and species. We analysed the computational performance of SMGen by generating batches of symmetric and asymmetric Reaction-based Models (RBMs) of increasing size, showing how a different number of reactions and/or species affects the generation time. Our results show that when the number of reactions is higher than the number of species, SMGen has to identify and correct high numbers of errors during the creation process of the RBMs, a circumstance that increases the overall running time. Still, SMGen can create synthetic models with 512 species and reactions in less than 7 seconds. The open-source code of SMGen is available on GitLab: https://gitlab.com/sgr34/smgen.

## 1. Introduction

Systems Biology is a multidisciplinary research field that combines mathematical, computational, and experimental expertise to understand and predict the behavior of complex biological systems [1,2]. Among the different formalisms that can be used to describe intracellular processes, Reaction-based Models (RBMs) [3–5] are the most suitable for obtaining a detailed comprehension of the mechanisms that control the emergent behavior of the system under analysis [4]. The analysis of RBMs can be used to drive the design of focused lab experiments; to this aim, computational tasks such as Parameter Estimation (PE), Sensitivity Analysis (SA), and Parameter Sweep Analysis (PSA) are generally applied [1,5–7]. Unfortunately, these computational tasks require the execution of huge amounts of simulations, so that the capabilities of biochemical simulators running on Central Processing Units (CPUs) (see, e.g., [8–10]) can be easily overtaken. Thus, several simulators exploiting Graphics Processing Units (GPUs) have been lately introduced to reduce the running times (see, e.g., [11–19]).

A crucial point, whenever new simulators are designed and implemented, regards the evaluation of their computational performance and their efficiency in executing the aforementioned demanding tasks. In this context, RBMs represent a key means as they can be exploited to run both stochastic simulation algorithms and (deterministic) numerical integration methods. Although there are more than 1,000 models of real biochemical systems publicly available on BioModels [20,21], with some of them characterized by hundreds of species and reactions, the majority of these models do not follow the law of mass-action. In addition, these models do not have the structural characteristics (e.g., the number of species and the number of reactions) necessary to perform a fair comparison among the simulators under different scenarios. Though, only a limited number of RBMs is present in the literature (e.g., signal transduction pathways [22–24] or metabolic pathways [25]). Thus, the lack of detailed RBMs, especially those characterized by hundreds or thousands of reactions and molecular species, thus hampers the possibility of performing a thorough analysis of the performance of these simulators.

The computational performance of several GPU-powered tools have been already assessed using randomly generated synthetic RBMs [13,18,19]. However, only a few generators of biochemical models have been proposed so far, hindering the possibility of having a common and well-defined benchmarking approach. For instance, Komarov *et al.* [13,14] developed a tool to generate synthetic networks, which was then used to test the performance of their GPU-based simulators. Given the number of reactants, the type of reactions to be included in the RBM, and the total number of reactions, they generated synthetic RBMs by exploiting a hash table to avoid duplicates. The tool was then modified by randomly sampling the values of the initial concentrations of the species from a uniform distribution and the kinetic constants from a logarithmic distribution [18]. Another known and established model generator is the Reaction Mechanism Generator (RMG) [26], which was specifically developed to create synthetic chemical processes. RMG exploits an extensible set of 45 reaction families to generate elementary reactions from chemical species, while the reaction rates are estimated using a database of known rate rules and reaction templates. RMG relies on graphs to represent the chemical structures, and trees to represent thermodynamic and kinetic data. Due to its peculiarities, RMG was used to, e.g., automatically create kinetic models for the conversion of bio-oil to syngas through gasification [27]. Finally, other tools, such as Moleculizer [28], were introduced for the generation of reaction systems to obtain a deeper understanding of transduction networks.

Despite the efforts done to automatically define synthetic models, all these generators share a common drawback, that is, they have a limited flexibility and can generate only a restricted set of biochemical networks and processes. As a matter of fact, the existing methods generally do not perform any check on the generated models; while some of them can be only used to create a restricted set of models (e.g., RMG can generate only elementary reactions). Moreover, in the case of the approach proposed by Komarov *et al.*, even thought it can generate first- and second-order reactions, including degradation and production reactions—which are the most common reactions in real biochemical networks—the tool is not publicly available.

Considering the impelling necessity of defining a common benchmarking approach that allows for fairly evaluating and comparing different simulation approaches [29], we propose here a novel tool, named SMGen, designed to automatically generate synthetic biological networks codified as RBMs, which allow to obtain non trivial dynamics. It is worth mentioning that we are mainly interested in the generation of RBMs suitable for the analysis and comparison of the performance of the biochemical simulators. An example of such RMBs is a model composed of plausible reactions (e.g., transformation, production and degradation reactions, which are common in real biochemical networks) leading to non trivial dynamics. For instance, in the case of a dynamics that instantly exhaust all reactants, some of the most advanced integration algorithms are able to simulate such stable and/or flat dynamics in just one computation step, thus hampering the possibility of a fair comparison among the different simulation approaches.

In addition, SMGen overcomes the existing approaches and tools, which generally do not allow for using different probability distributions for the initialization of the species amounts and the kinetic constants. As a matter of fact, the possibility of exploiting different probability distributions for the initialization of these parameters is an important features offered by SMGen, since the effort required for the simulation of the same model can vary according to the model parameterizations (i.e., species amounts and the kinetic constants).

SMGen adheres to well-defined structural characteristics based on graph theory and linear algebra properties, in particular, it exploits the definition of an undirected graph with a single connected component, which makes the whole generation process a computationally demanding task. To overcome this limitation, on the one hand, SMGen internally codifies all data structures by means of sparse matrices as well as *ad-hoc* structures specifically designed to avoid worthless values that would increase the running time required to generate RBMs. On the other hand, SMGen is able to drastically reduce the computational time by exploiting a Main-Worker paradigm used to distribute the overall generation process of RBMs onto multi-core CPUs. We show that SMGen can create, in less than 7 seconds, synthetic RBMs with hundreds of chemical species and molecular reactions, whose dynamics exhibit non trivial characteristics of real biochemical networks. Among the different features provided by SMGen, it allows for easily generating both symmetric and asymmetric RBMs: symmetric RBMs are composed of a number of species equal to the number of reactions, while in asymmetric RBMs the number of species can be lower than the number of reactions or vice-versa. From a computational point of view, the concept of symmetry is crucial in the analysis of complex networks to measure their information and entropy [30]. In addition, considering that every RBM can be converted into the corresponding system of coupled Ordinary Differential Equations (ODEs), studying the symmetries of this system of ODEs can reveal the intrinsic properties of the system of interest [31]. As a matter of fact, Ohlsson *et al.* pointed out that an alternative analysis of the system of ODEs can be carried out by considering the symmetries of the system solutions, aiming at formalizing the structures and behavior of the underlying dynamics of biological systems. Moreover, the possibility of evaluating GPU-powered simulators using symmetric and asymmetric RBMs is fundamental to understand their performance under different conditions. Indeed, a fair comparison would allow the user to select the best simulator based on characteristics of the RBM that has to be analysed. Thanks to its features and efficiency, SMGen was used to generate the synthetic RBMs necessary to realize a thorough comparison of the performance of different meta-heuristics in solving the PE problem of biochemical networks [3,5], which is one of the most common and difficult computational issues in Systems Biology. The outcome of such analyses showed that some well-known and widely used meta-heuristics, generally able to outperform all competitors in the optimization of benchmark functions, obtained poor performance when used to solve the PE problem. Moreover, SMGen was exploited to generate symmetric and asymmetric RBMs required for the in-depth analyses and comparisons of the computational performance of different biochemical simulators and to highlight their peculiarities [19]. In particular, these analyses allowed for determining the best simulator to employ under specific conditions, such as the size of the RBM and the total number of simulations to perform. SMGen can also be used for the generation of biochemical networks to investigate the performance of approaches specifically designed to tackle other well-known and computationally demanding tasks in Systems Biology (e.g., SA and PSA [1,7]).

SMGen allows also for exporting the generated RBMs into the Systems Biology Markup Language (SBML) [32], Version 4 Level 2, and into the BioSimWare standard [33], which is used by different GPU-powered simulators. Thus, we designed and developed SMGen to be a unifying, user-friendly, and standalone tool freely accessible to the Systems Biology community. The RBMs can be easily generated by using the provided user-friendly Graphical User Interface (GUI), which is designed to help the users in setting all the parameters required to generate the desired RBMs.

The manuscript is structured as follows. Section 2 describes the mathematical formalism of RBMs, as well as the structural characteristics that must be complied to generate synthetic biological networks. In addition, we provide all the algorithms and details at the basis of SMGen. Section 3 shows the experimental results achieved by SMGen. Finally, a discussion and conclusive remarks are provided in Section 4.

## 2. Materials and Methods

### 2.1. Reaction-based Models

An RBM is defined by specifying the set 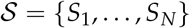 molecular species, and the set 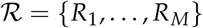 biochemical reactions that describe the interactions among the species appearing in 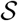. Each reaction *R_i_*, with *i* = 1,…, *M*, is defined as:

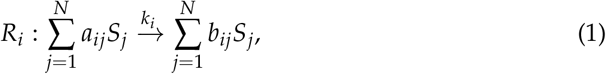

where *a_ij_* and 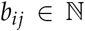 are the stoichiometric coefficients, and *k_i_* ∈ ℝ^+^ is the kinetic constant associated with *R_i_*. The stoichiometric coefficients specify how many molecules of species *S_j_*, with *j* = 1,…, *N*, appear either as reactants or products in reaction *R_i_*. Note that some species might not appear in a reaction, so that the corresponding stoichiometric coefficient will be equal to 0. The order of a reaction is equal to the total number of molecules (of the same or different species) that appear as reactants in that reaction.

Each RBM can be written in the compact matrix-vector form 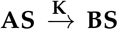, where **S** = [*S*_1_ ⋯ *S_N_*]^⊤^ is the *N*-dimensional column vector of the molecular species, **K** = [*k*_1_ ⋯ *k_M_*]^⊤^ is the *M*-dimensional column vector of the kinetic constants, while 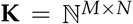 are the stoichiometric matrices, whose non-negative elements [*A*]_*i,j*_ and [*B*]_*i,j*_ correspond to the stoichiometric coefficients *a_ij_* and *b_ij_* of the reactants and products of the reactions, respectively.

Starting from an RBM and assuming the law of mass-action [34–36], the system of coupled ODEs corresponding to the RBM can be derived as follows:

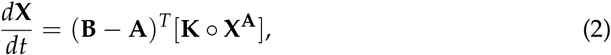

where each ODE describes the variation in time of a species’ concentration. In Equation 2 the *N*-dimensional vector **X** = [*X*_1_ ⋯ *X_N_*] represents the concentration values of species *S*_1_,…, *S_N_*, while **X^A^** is the vector-matrix exponentiation form [34]; the symbol ○ denotes the entry-by-entry matrix multiplication (Hadamard product).

### 2.2. SMGen

In order to generate synthetic models of biochemical networks, SMGen complies with specific structural characteristics that the RBMs have to satisfy, that is:

- *System connectivity:* a biochemical network can be represented as an undirected graph with a single connected component, where the nodes represent the molecular species and the edges correspond to the species interactions. This representation easily allows for ensuring the system connectivity, a property that is strictly required to ensure that each species 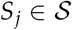, with *j* = 1,…, *N*, will be involved in at least one reaction 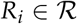, with *i* = 1,…, *M*. To be more precise, as a first step, an undirected graph with a single connected component is built. This undirected graph is randomly generated by using *N* – 1 edges and by taking into account the maximum number of reactants and products, obtaining a connected biochemical reaction network. It is worth noting that a graph with a single connected component can be built if and only if the following condition is met:

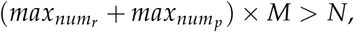

where *max_num_r__* and *max_num_p__* are the maximum number of reactants and products, respectively. As a second step, starting from the initial undirected graph, the stoichiometric matrices, which can be viewed and treated as a Petri net [37,38] that, in turn, can be considered as a bipartite graph, are generated. Then, the stoichiometric matrices are randomly updated by adding and removing connections among the species, always taking into consideration the maximum number of reactants and products. Note that the initial connections, which correspond to the edges of the initial undirected graph, are never removed to ensure that all the species are involved in at least a reaction, maintaining the whole biochemical network connected. We designed an algorithm (see Algorithm 2) that builds the graph, composed of a single connected component, with the minimum number of edges that are needed to connect all the nodes.
- *Maximum number of reactants and products:* for each reaction 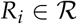, with *i* = 1,…, *M*, the number of reactants and the number of products cannot be arbitrarily large, but has to be lower than or equal to a user-defined values (i.e., *max_num_r__* and *max_num_p__*). Stated otherwise, the maximum order of the generated reactions should be fixed. It is worth mentioning that SMGen does not explicitly account for conservation conditions during the generation of the RBMs. However, it can generate reactions that are akin, for example, to protein modifications or conformational changes, thus resulting in two biochemical species whose sum is constant during the simulation.
- *Linear independence:* to ensure that each reaction *R_i_*, with *i* = 1,…, *M*, is endowed with plausible characteristics of a real biochemical reaction, the vectors of the stoichiometric coefficients of the reactants and products involved in *R_i_* must be linearly independent. The linear independence condition allows for avoiding unrealistic reactions. An example of an unrealistic reaction consists in increasing or decreasing the amount of a species starting from one molecule of the species itself, e.g., 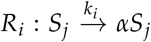, with *α* > 1. Since the linear independence is evaluated between the reactants and the products of each reaction, it is related to the number of species and reactions. Thus, it could happen that when the number of species is much higher than the number of reactions it is not possible to build a graph with a single connected component. In such a case, duplicated reactions or linear dependent vectors between reactants and products of some reactions will occur. On the contrary, it is always possible to build a graph, composed of a single connected component, when the number of reactions is much higher than the number of species.
- *Reaction discreteness:* each reaction *R_i_*, with *i* = 1,…, *M*, must appear only once in the network, that is, duplicated reactions are not allowed.

SMGen is provided with a user-friendly GUI (see Figure 1) that allows the user to easily set up all the parameters required to generate the desired synthetic RBMs:

- the number of species *N* and the number of reactions *M*;
- the maximum number of reactants and products *max_num_r__* and *max_num_p__* that might appear in any reaction;
- the probability distribution 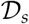 that is used to initialize the species amounts (to be chosen among uniform, normal, logarithmic or log-normal distributions). It is worth mentioning that all species amounts are initialized using the same distribution probability;
- the minimum and maximum values *min_s_* and *max_s_* for the initial species amounts (to be specified either as number of molecules or concentrations);
- the probability distribution 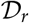 that is used to set the values of the kinetic constants (to be chosen among uniform, normal, logarithmic or log-normal distributions). Note that all kinetic constants are generated using the same distribution probability;
- the minimum and maximum values *min_r_* and *max_r_* for the kinetic constants;
- the total number of RBMs that the user wants to generate;
- the output format file to export the generated RBMs (i.e., BioSimWare [33] and SBML [32]);
- the mean and standard deviation values *μ_s_* and *σ_s_* for the initial amounts—as well as the mean and standard deviation values *μ_r_* and *σ_r_* for the kinetic constants—must also be provided if the normal or log-normal distributions are selected.

**Figure 1.**
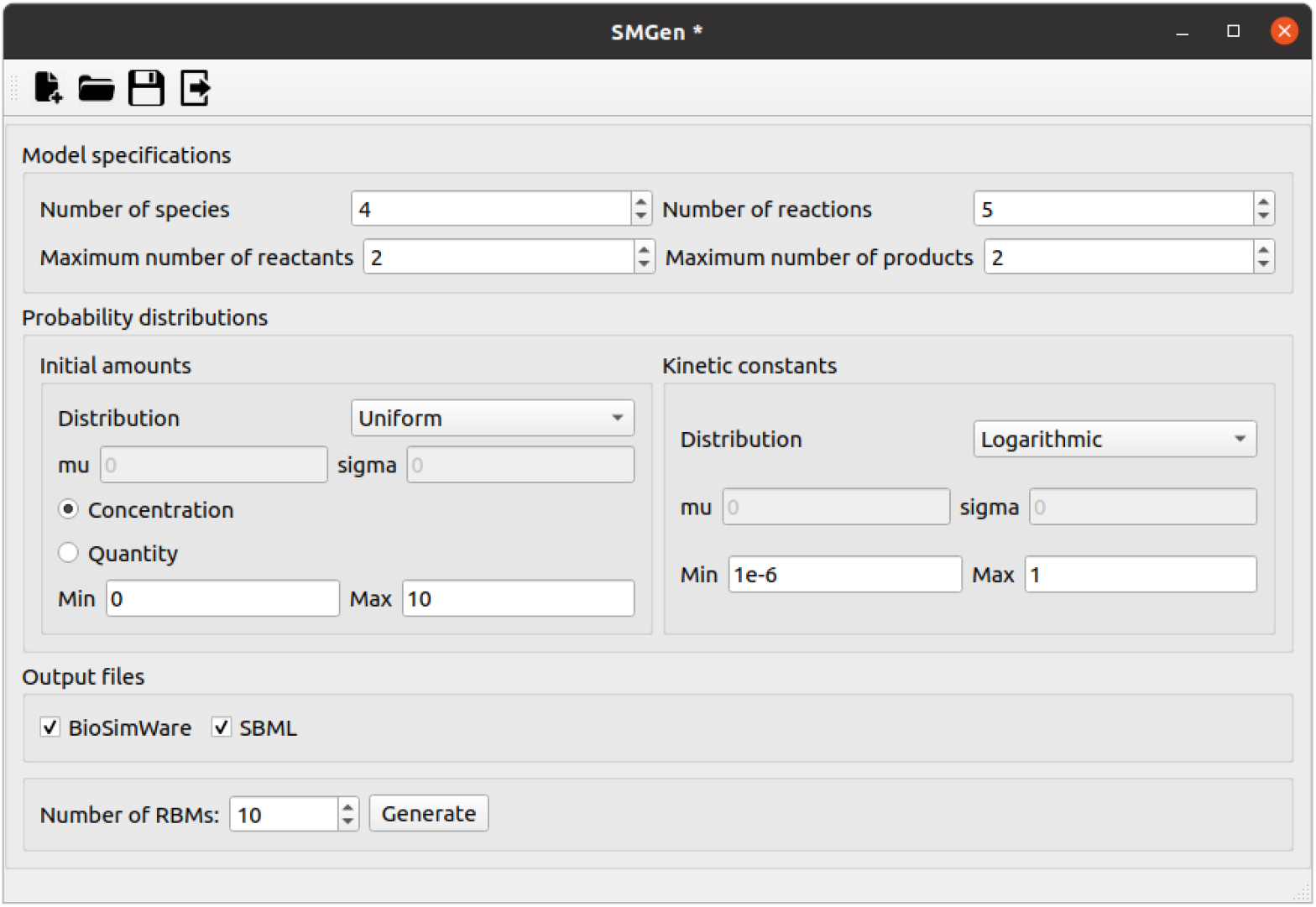
Graphical User Interface of SMGen. The user can set all the parameters to generate the desired RBMs, i.e., number of species and reactions, maximum number of reactants and products, probability distribution for the initial amounts and kinetic constants, and the output format file (i.e., BioSimWare, SBML).

Figure 2 shows a high-level scheme of the proposed implementation of SMGen, which exploits the Main-Worker paradigm to speed up the generation of the RBMs [39]. The user can specify the number of processes *P*—otherwise automatically set to the minimum value 3—which are used as follows:

- Proc_1_ manages the GUI;
- Proc_2_ is the Main process that orchestrates the computation;
- Proc_*p*_, with *p* = 3,…, *P*, are the Worker processes. The whole functioning of SMGen can be summarized as follows:
- the user interacts with the GUI, managed by Proc_1_, to fill in all the required values for the parameters necessary to create the RBMs;
- Proc_1_ sends the values of all parameters to the Main process (Proc_2_), which allocates the resources and distributes the work to the Workers (Proc_*p*_, with *p* = 3,…, *P*);
- each Worker (Proc_*p*_, with *p* = 3,…, *P*) generates an RBM. As soon as a Worker terminates its execution, it communicates to the Main process that the RBM has been created. If necessary, the Main process assigns the generation of other RBMs to idle Workers. When all required RBMs are obtained, the Workers enter in the death state, while the Main process waits for further instructions from Proc_1_.

**Figure 2.**
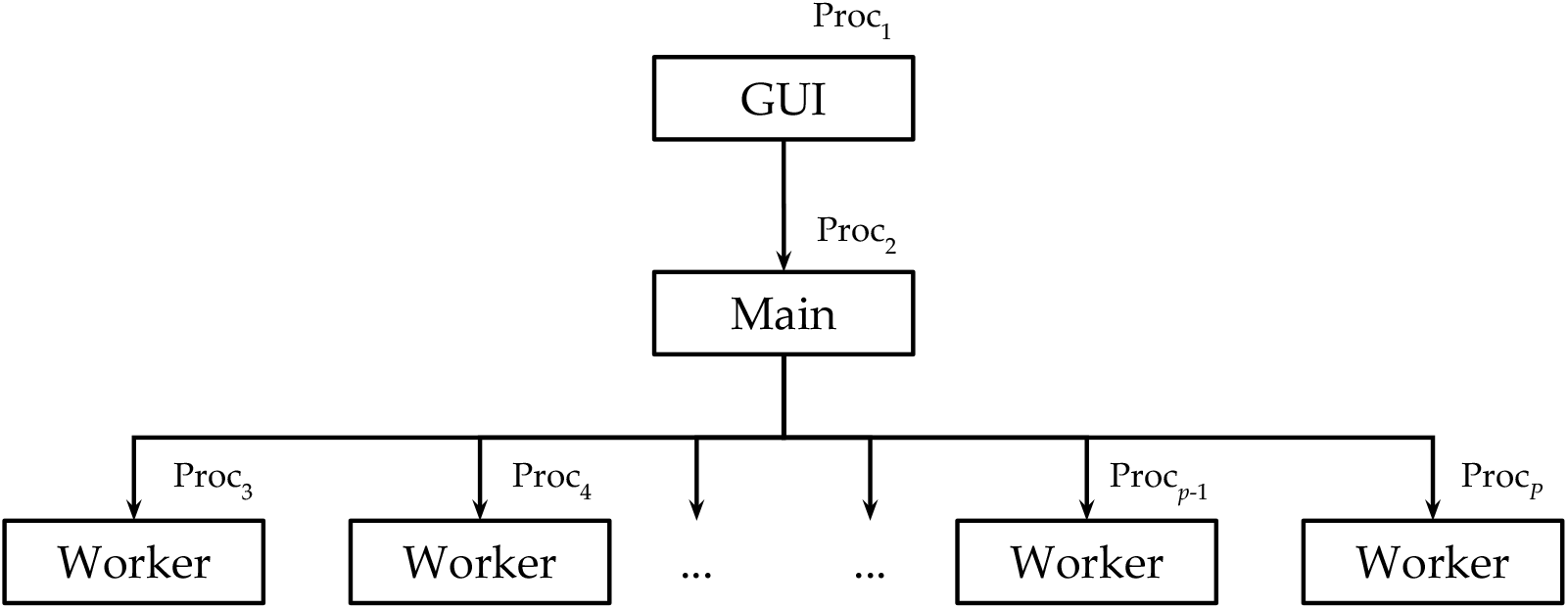
Scheme of the Main-Worker implementation of SMGen. The Main process (Proc_2_) orchestrates all the available Workers (Proc_*p*_, with *p* = 3,…, *P*), which generate the RBMs in a distributed computing fashion.

**Figure 3.**
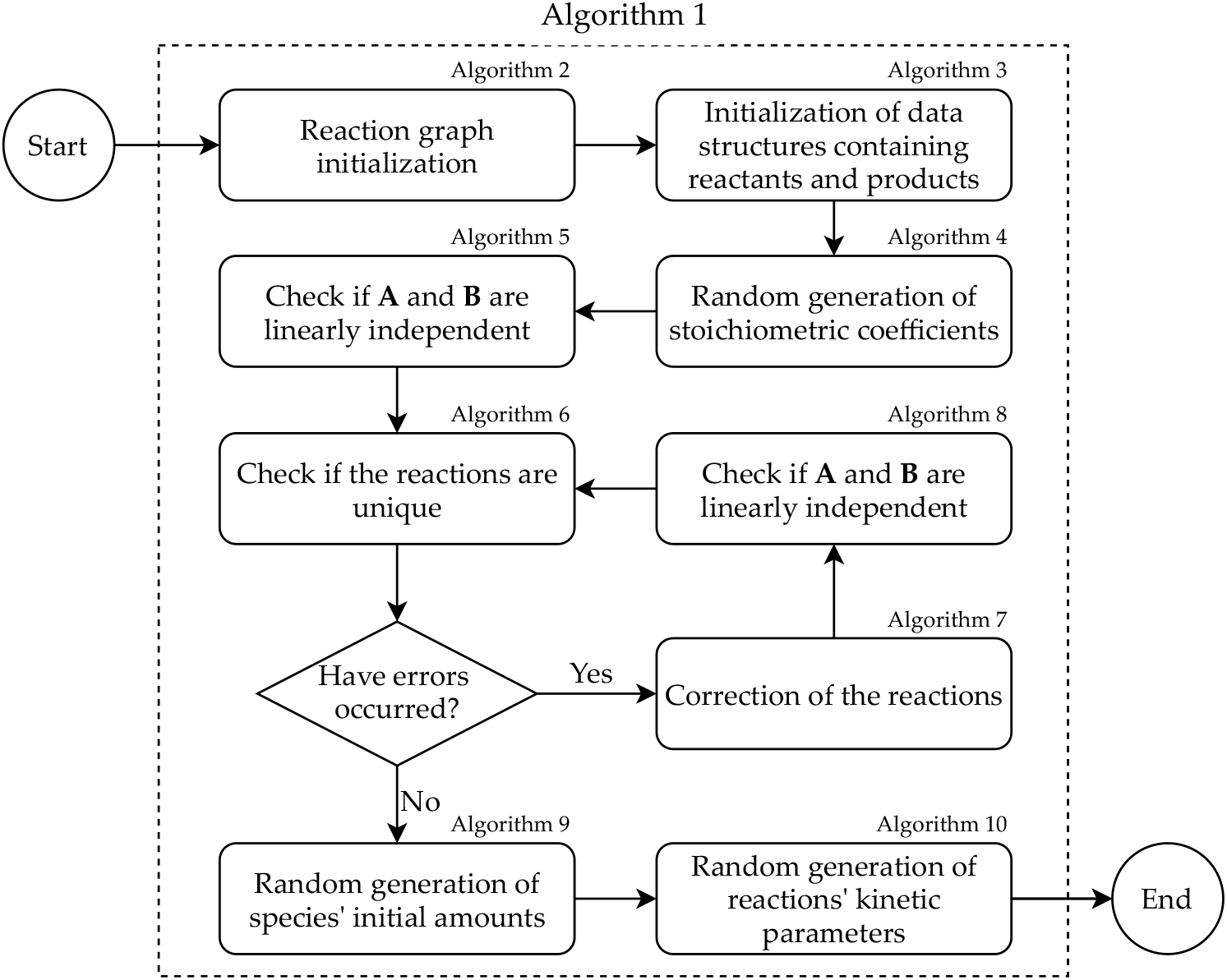
Workflow of a single Worker execution. First, the graph of the reactions is randomly initialized, and then converted into the data structures used to store the reactants and products. Second, the stoichiometric coefficients are randomly generated and the consistency of the reactants and products is verified. Third, the initial amounts of the species and the kinetic parameters of the reactions are randomly generated using the probability distributions specified by the user.

The workflow of each Worker consists in 9 different phases, in which a specific algorithm is executed (see Figure 3).

**Algorithm 1.**
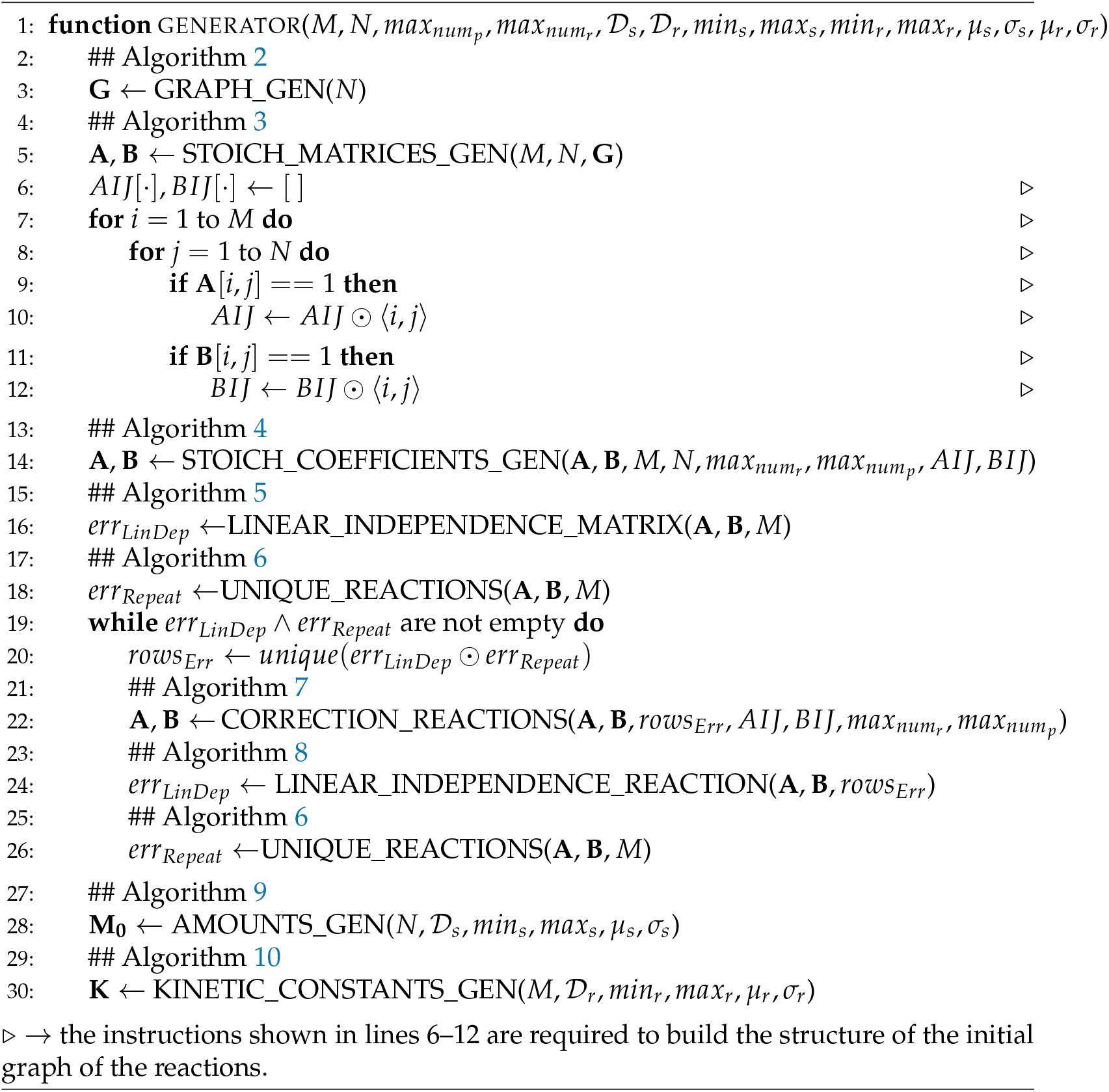
SMGen: workflow of a single Worker execution.

The pseudo-code reported in Algorithm 1 briefly summarizes all the steps required to generate a single RBM; the pseudo-code of the procedures invoked within Algorithm 1 are reported in Appendix A. For the sake of clarity, Table 1 lists the symbols used in the following description and in the pseudo-codes.

**Table 1:** List of symbols used in the pseudo-code of algorithms at the basis of SMGen.

| Symbol | Description |
| --- | --- |
| $M$ | Number of reactions composing the RBM |
| $N$ | Number of species involved in the RBM |
| $max_{num_r}$ | Maximum number of the reactants |
| $max_{num_p}$ | Maximum number of the products |
| $\mathbf{M}_0$ | Array of the initial amounts |
| $\mathbf{K}$ | Array of the kinetic constants |
| $\mathbf{A}$ | Stoichiometric matrix of the reagents |
| $\mathbf{B}$ | Stoichiometric matrix of the products |
| $\mathbf{G}$ | Adjacency matrix of the graph of the reactions |
| $\mathcal{D}_s$ | Probability distribution for the initial amounts |
| $min_s$ | Minimum value of the initial amounts |
| $max_s$ | Maximum value of the initial amounts |
| $\mu_s$ | Mean of the normal and log-normal distributions for the initial amounts |
| $\sigma_s$ | Standard deviation of the normal and log-normal distributions for the initial amounts |
| $\mathcal{D}_r$ | Probability distribution for the kinetic constants |
| $min_r$ | Minimum value of the kinetic constants |
| $max_r$ | Maximum value of the kinetic constants |
| $\mu_r$ | Mean of the normal and log-normal distributions for the initial amounts |
| $\sigma_r$ | Standard deviation of the normal and log-normal distributions for the kinetic constants amounts |
| $\odot$ | The concatenation operator |

The steps performed by each Worker to generate an RBM are the following:

1. Given the parameters provided by the user, the graph representing the species and their interactions is randomly initialized (line 3 of Algorithm 1, see Algorithm 2).
2. The adjacency matrix of the graph generated in Step 1 is converted into the stoichiometric matrices **A** and **B** (line 5 of Algorithm 1, see Algorithm 3). Note that the instructions in lines 6–17 of Algorithm 1 are required to build the data structure of the initial graph, which is then modified.
3. The stoichiometric coefficients are randomly generated (line 19 of Algorithm 1, see Algorithm 4).
4. For each reaction *R_j_*, with *i* = 1,…, *M*, the linear independence between the reactants and products is verified (line 21 of Algorithm 1, see Algorithm 5).
5. The uniqueness of each reaction in the RBM is verified (line 23 of Algorithm 1, see Algorithm 6).
6. Any error in the RBM identified in the previous steps is corrected (line 27 of Algorithm 1, see Algorithm 7); the linear independence and the uniqueness of the reactions in the modified RBM are iteratively verified (lines 29 and 31 of Algorithm 1, see Algorithms 8 and 6, respectively).
7. The initial amounts of the species are generated according to the chosen probability distribution (line 33 of Algorithm 1, see Algorithm 9). If a species appears only as a reactant in the whole RBM, its amount is set to remain unaltered. The rationale behind this is double: on the one hand, we avoid the possibility of creating reactions that could be applied at most once, which is a highly improbable situation in biological systems; on the other hand, we mimic the non-limiting availability of some biochemical resources, for instance, it might be used to reproduce the execution of *in vitro* experiments where some species are continually introduced in the systems to keep their amount constant [40].
8. The kinetic constants of the reactions are generated according to the chosen probability distribution (line 35 of Algorithm 1, see Algorithm 10).

SMGen was developed using the Python programming language and exploiting `mpi4py` [41], which provides bindings of the Message Passing Interface (MPI) specifications for Python to leverage multi-core CPUs [42]. The open-source code of SMGen is available on GitLab (https://gitlab.com/sgr34/smgen) under the GNU GPL-3 license. In addition, it can be easily installed using the Python package installer pip (https://pypi.org/project/smgenerator).

## 3. Results

We analyzed the performance of SMGen regarding both its capability of creating RBMs characterized by non trivial dynamics, and the computational time required to generate sets of RBMs of increasing size. All tests were executed on a workstation equipped with an Intel Core i7-8750H CPU (clock 4.1 GHz), 16 GB of RAM and a Samsung 970 EVO solid-state drive NVMe PCIe (up to 3400 MB/s and 1500 MB/s read and write speed, respectively), running on Ubuntu 20.04 LTS.

As a first batch of tests, we generated 100 synthetic RBMs characterized by a limited number of reactions and species (4 and 5, respectively), and we analysed their characteristics and dynamics. We set to 3 both the maximum number of reactants *max_num_r__* and products *max_num_p__*. We sampled the initial amounts of species from a normal distribution with mean *μ_s_* = 5 and standard deviation *σ_s_* = 5, considering a minimum value *min_s_* = 0 and maximum value *max_s_* = 10. The kinetic constants were instead sampled from a logarithmic distribution with minimum value *minr* = 10^−16^ and maximum value *max_r_* = 10.

Table 2 shows the list of reactions along with the kinetic constants of one of these 100 synthetic RBMs. Since the species *X*_0_ appears only as a reactant, its amount will be kept constant during the simulation. The initial molecular amounts of all species—given as number of molecules—are listed in Table 3. This small RBM includes the basic “cascade of reactions” structure typically observed in signaling pathways, starting from the source represented by species *X*_0_ and *X*_4_, toward species *X*_2_ and *X*_3_.

**Table 2:**
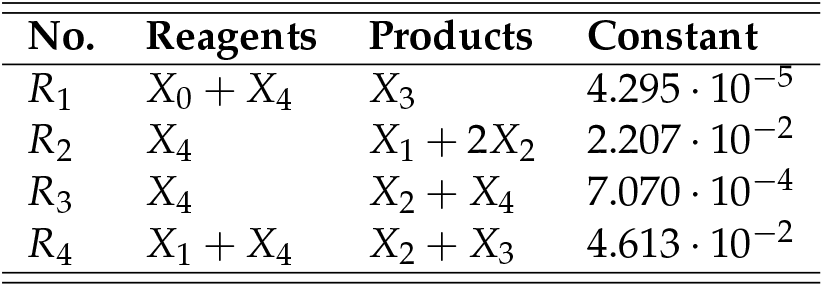
List of the reactions of an RBM with by 4 reactions and 5 species generated by SMGen.

| No. | Reagents | Products | Constant |
| --- | --- | --- | --- |
| $R_1$ | $X_0 + X_4$ | $X_3$ | $4.295 \cdot 10^{-5}$ |
| $R_2$ | $X_4$ | $X_1 + 2X_2$ | $2.207 \cdot 10^{-2}$ |
| $R_3$ | $X_4$ | $X_2 + X_4$ | $7.070 \cdot 10^{-4}$ |
| $R_4$ | $X_1 + X_4$ | $X_2 + X_3$ | $4.613 \cdot 10^{-2}$ |

**Table 3:** Initial molecular amounts of the RBM generated by SMGen shown in Table 2.

| Species | Initial amount |
| --- | --- |
| $X_0$ | 4 |
| $X_1$ | 8 |
| $X_2$ | 7 |
| $X_3$ | 8 |
| $X_4$ | 1 |

We simulated the dynamics of this RBM for 50 time steps (arbitrary units), and the achieved dynamics are shown in Figure 4. These plots evidence that, although the RBM was randomly generated by SMGen, the simulated behavior is non trivial (e.g., the reactants are not instantly exhausted, resulting in a flat dynamics). Note that currently SMGen can be only used to generate RBMs and does not include any simulation tools yet. The simulation of the dynamics of this RBM was carried out using FiCoS [19]. It is worth mentioning that obtaining synthetic RBMs exhibiting non trivial dynamics is fundamental to perform in-depth computational analyses and comparisons among the existing and the novel simulators. Indeed, in the case of stable or flat dynamics, or when the overall behavior of the network is extremely fast and instantly exhausts all the reactants, the most advanced integration algorithms are able to simulate the emergent dynamics in just one computation step [19]. In such a case, the computational performance of the simulation tools is only partially assessed, thus hindering a fair comparison among the tools.

**Figure 4.**
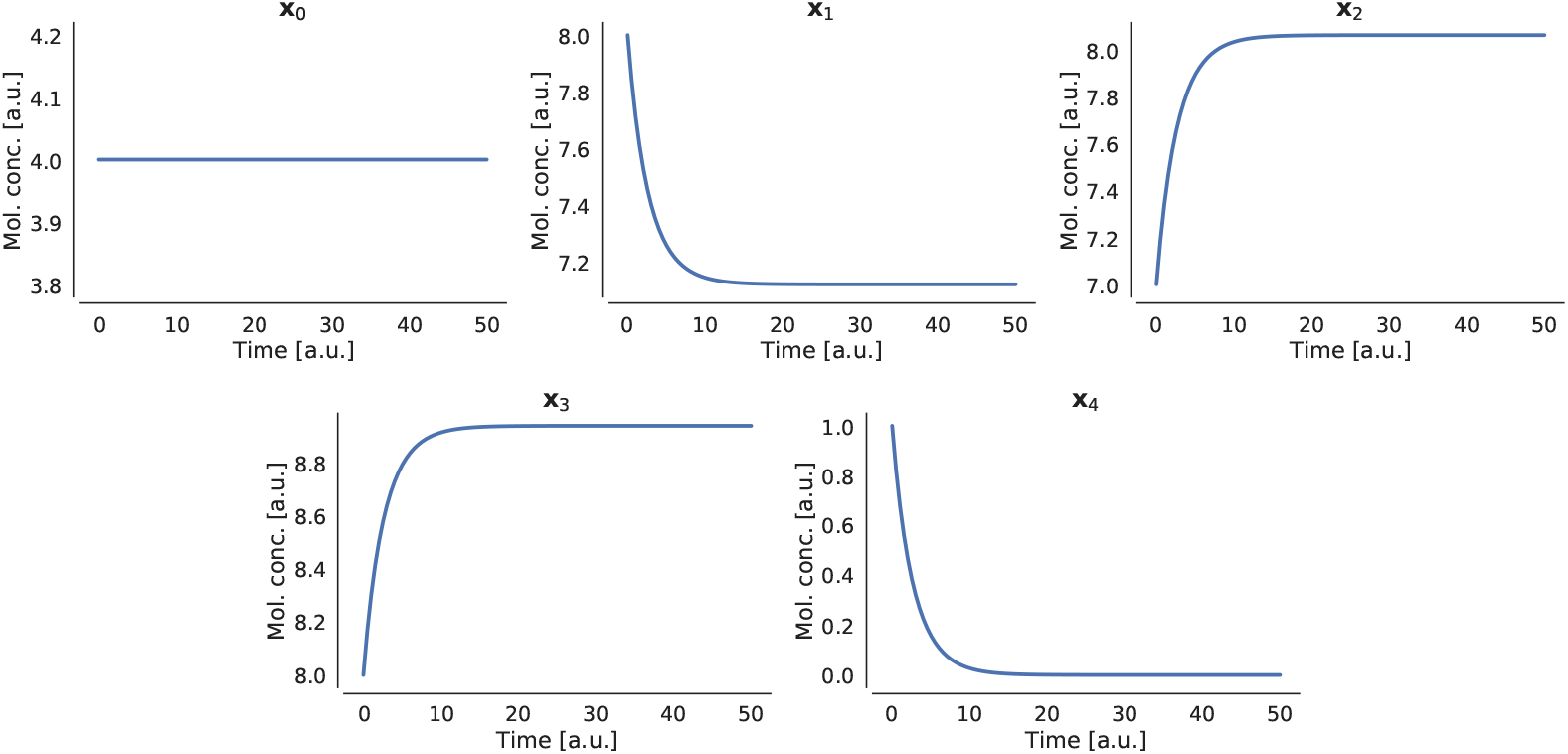
Dynamics of the species of the synthetic RBM generated by SMGen shown in Table 2.

As a second batch of tests, we evaluated the computational performance of SMGen exploiting the Main-Worker paradigm running on 4 distinct cores of the CPU. First, we considered the generation of symmetric RBMs with an increasing number of species and reactions (i.e., *M* = *N* = 2^*x*^, with *x* = 2,…,9). The initial amounts and kinetic constants were randomly sampled from a uniform distribution with minimum values *min_s_ = min_r_* = 0 and maximum values *max_s_* = *max_r_* = 10. We also varied the maximum numbers of reactants and products considering the set of values {2,3,4} and setting *max_numr_ = max_num_p__*. For each of the resulting 24 parameters combinations, we created 100 RBMs to collect statistically sound results about the performance of SMGen. As described in Section 2, two kinds of error can occur during the generation of an RBM: a linear dependence between reactants and products, and duplicated reactions. Since the correction of these errors is one of the most time-consuming phases of SMGen, we separately measured the generation time, which indicates the running time spent by SMGen to generate an RBM, and the saving time, which refers to the writing operations on the solid-state drive. Figure 5 shows the average running time required by SMGen to generate and save an RBM. As expected, both the generation and saving time increase along with the number of species and reactions of the RBM. Moreover, we observe that the maximum number of reactants and products have a slight impact on both the generation and the saving time; in most of the cases, increasing these values results in a higher running time.

**Figure 5.**
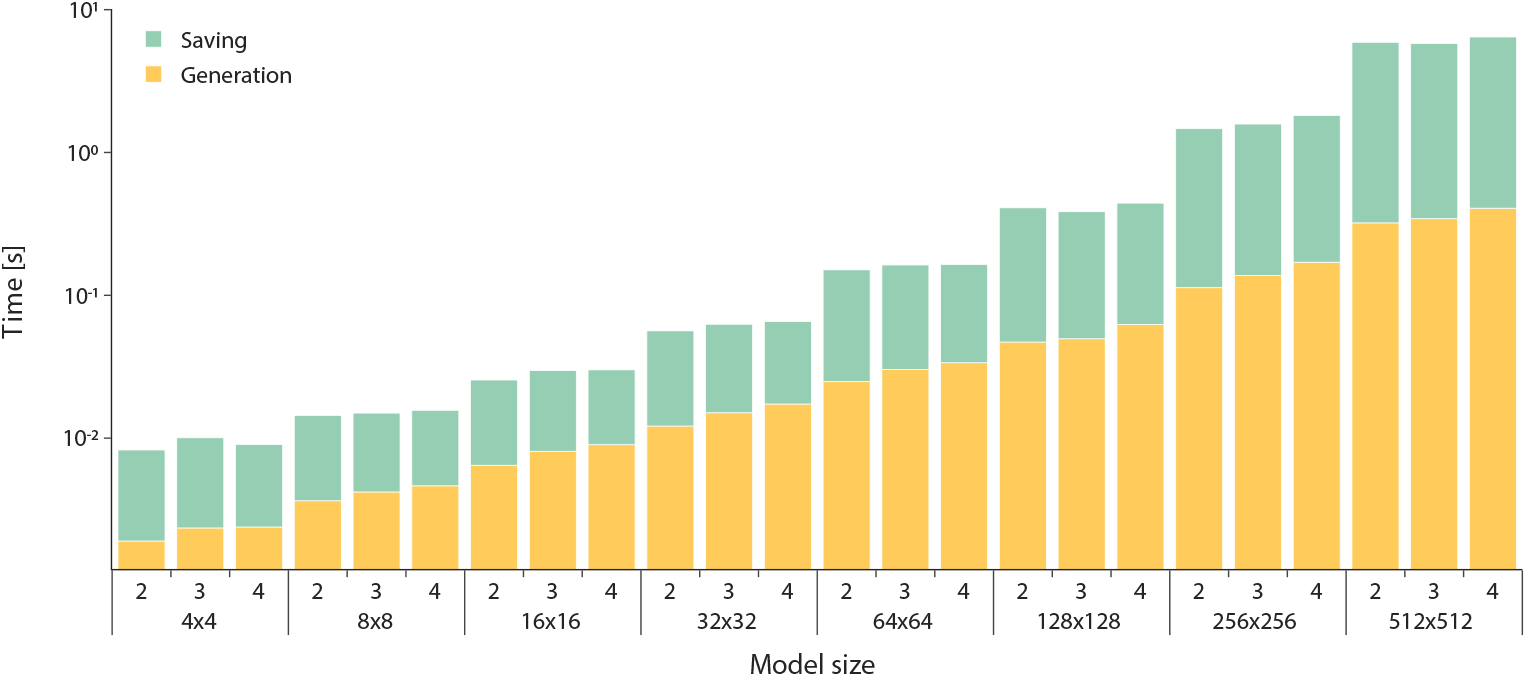
Stacked bar plot showing the average generation time (yellow bars) and the average saving time (green bars) required by SMGen to generate a symmetric RBM. Note that the y-axis is in logarithmic scale.

Finally, we exploited SMGen for the creation of asymmetric RBMs, to evaluate how a different number of species and reactions affects the running time. As in the case of symmetric RBMs, we measured both the generation time and the saving time. The asymmetric RBMs were created as follows:

- we set the number of species *N* ∈ {4,8,16,32,64}, and then we varied the number of reactions *M* ∈ {2*N*,4*N*,8*N*};
- we set the number of reactions *M* ∈ {4,8,16,32,64}, and then we varied the number of species *N* ∈ {2*M*,4*M*,8*M*};
- we varied both the maximum numbers of reactants *max_num_r__* and products *max_num_p__* in {2,3,4}.

**Figure 6.**
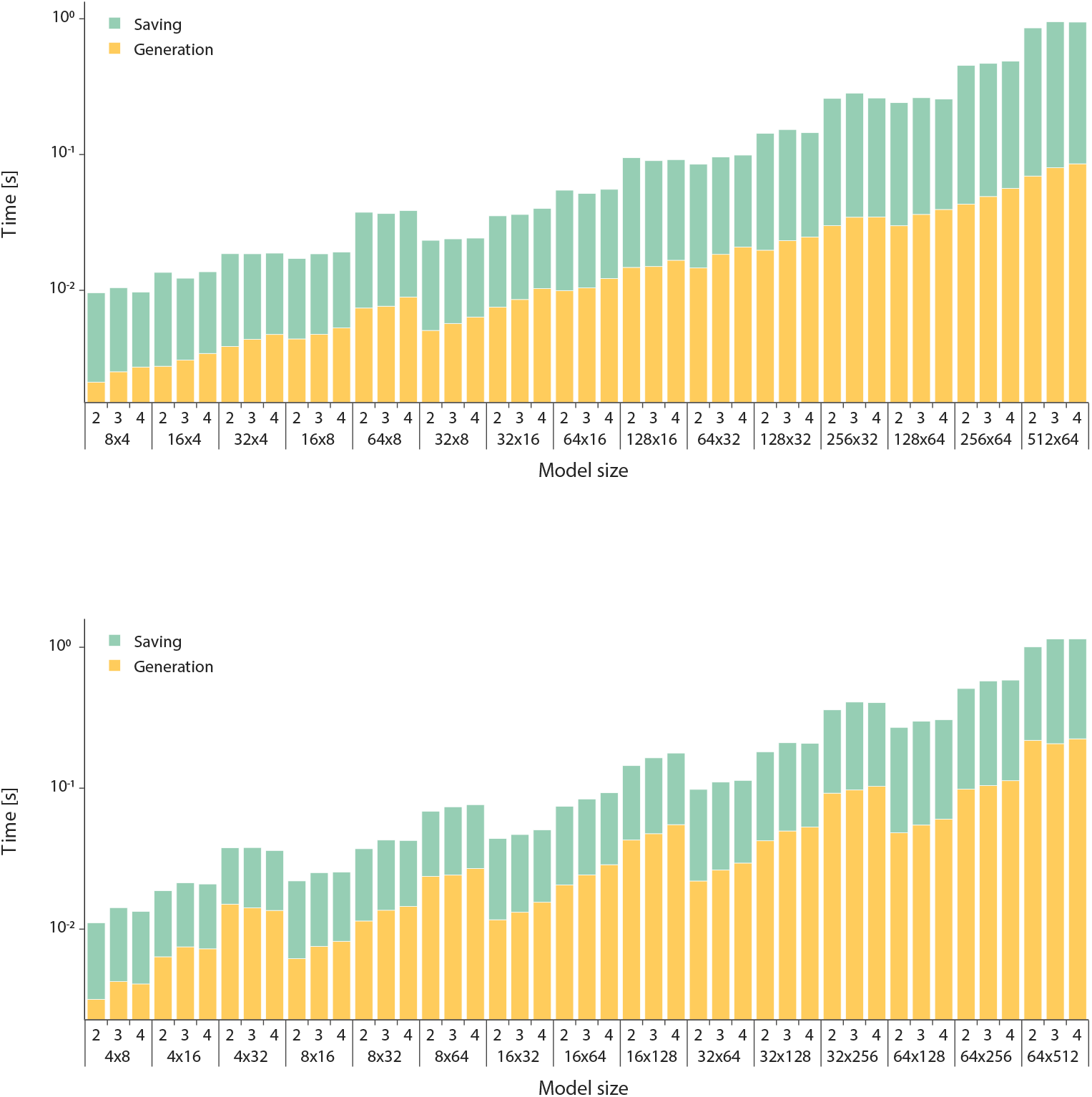
Stacked bar plots showing the average generation time (yellow bars) and the average saving time (green bars) required by SMGen to generate an asymmetric RBM with more species than reactions (top) and with more reactions than species (bottom). Note that the *y*-axes are in logarithmic scale.

In such a way, we obtained a total of 90 different combinations of the parameters (i.e., number of species, number of reactions, and maximum number of reactants and products) to be tested; as in the previous tests, for each combination we generated 100 RBMs to collect statistically sound results. Figure 6 shows the average running time required to create RBMs with dimensions *N* × *M*, highlighting once again that both the generation time and the saving time increase along with the size of the RBMs. As in the case of symmetric RBMs, we observed the same effect due to the maximum number of reactants and products allowed in the reactions. As expected, when there are more reactions than species (bottom panel in Figure 6) the generation times are higher than the opposite situation (top panel in Figure 6). This circumstance is due to the potential higher number of errors that SMGen has to identify and correct. Indeed, when *M* ≫ *N*, the probability that repeated reactions are randomly generated is higher than the case when *N* ≫ *M*, because the number of admissible reactions strictly depends on the number of species.

## 4. Conclusions

In this work we presented SMGen, a generator of synthetic RBMs displaying the characteristics of real biochemical networks, which can be exploited to create benchmarks for the evaluation of novel and existing simulators. SMGen is particularly suitable to create the RBMs necessary to assess the performance of GPU-based simulators. As a matter of fact, the performance of GPU-powered simulators can drastically change with the number of chemical species and reactions composing an RBM. Considering that every RBM can be converted into the corresponding system of coupled ODEs, the resolution of this system of ODEs can be performed in a parallel fashion, where each ODE is resolved by a thread. Since each ODE corresponds to a specific chemical species, a higher number of species generally lead to a higher parallelization, increasing the computational performance of the simulator. On the contrary, considering that the number of the reactions composing the biological system is roughly related to the length of each ODE, in terms of the mathematical complexity, the higher the number of reactions the higher the number of operations that must be performed by each thread, leading to a higher running time [18,19].

SMGen was developed in Python and was designed to be a unifying, user-friendly, and standalone tool. In addition, SMGen exploits the Main-Worker paradigm to speed up the generation of RBMs; this was implemented using the mpi4py [41] library, where the first process manages the GUI, the second one is the Main process, and all the other processes are the Workers that generate the RBMs in a distributed computing fashion. Thanks to the GUI of SMGen, the user can easily set up all the parameters characterizing the required RBMs, e.g., the number of species and reactions, the maximum number of reactants and products per reaction, the probability distributions (uniform, normal, logarithmic, log-normal) to generate the initial amounts of the species and the values of the kinetic constants associated with the reactions, the output file format to save the RBMs. It is noteworthy that the performance of SMGen is not affected by the choice of different distributions for the sampling of the species amounts and kinetic constants. On the contrary, different distributions can drastically affect the dynamics of the model (see, e.g., the Brusselator model [38,43]). The analysis of the features of the generated models, according to different distributions, will be addressed in a future work.

We assessed the capabilities of SMGen for the creation of RBMs characterized by a non trivial behavior, and we presented an example of a synthetic RBM together with the simulated dynamics. We also tested the computational performance of SMGen by generating batches of symmetric and asymmetric RBMs of increasing size, showing the impact of the number of reactions and species, and of the number of reactants and products per reaction, on the generation times. We observed that when the number of reactions is higher than the number of species, SMGen generally identifies and corrects high numbers of errors during the creation process of the RBMs, a circumstance that inevitably increases the overall running time.

As a future extension of this work, we plan to develop an Application Programming Interface (API), so that SMGen can be seamlessly integrated into other processing pipelines, tools, and simulators. We are also developing a well-documented commandline interface, which will be released together with the API, to increase the user experience as well as the integrability with the existing approaches. Another feature that could be introduced is a module for the visualization of the networks. However, even though the visualization of the networks might help the users, plotting RBMs composed of thousands of species and reactions is quite a hard task. Moreover, the visualization of such large RBMs might not add any relevant information. We will also introduce a new feature specifically developed to generate feedback loops in synthetic RBMs, exploiting the theory of Petri nets [37,38]. Feedback loops are fundamental elements of biological processes that lead to the establishment of oscillatory regimes and non-linear dynamics [22]. Moreover, we plan to develop a function to rescale the kinetic constants by taking into account the number of reactants involved in the reactions. As a matter of fact, both the order of magnitude and the value of each kinetic constant are related to the number of reactants composing the corresponding reaction. In the current version of SMGen, we assume that all reactions follow the law of mass-action; however, we will implement additional kinetics (e.g., Michaelis-Menten and Hill kinetics [44,45]) in the future.

SMGen relies on graph theory and linear algebra properties to comply with specific structural characteristics that are essential to create synthetic but real RBMs. Nevertheless, the current version of SMGen does not exploit network graphlets and motifs, which characterize families of real biological networks. Graphlets are induced sub-graphs appearing at any frequency because they are independent of the network’s null model (i.e., a random graph model defined by a probability distribution), while network motifs are repeated sub-graphs appearing with a frequency higher than in random graphs and depending on the network’s null model [46]. Graphlets have been used to analyze local network structures, and to cluster different network types [46]. Probabilistic graphlets have been also developed to analyze the local wiring patterns of probabilistic networks. When applied to study biological networks, probabilistic graphlets resulted in a robust tool, able to capture the biological information underlying the biological networks, thanks to the capabilities of managing the low signal topology information [47]. Graphlets can also be used to measure the structural similarity among large networks, calculating the graphlet degree distribution [48]. Motifs can be used to study autoregulation, single input module, dense overlapping regulons and feedback loops, as well as to reveal answers to many important biological questions [49,50]. The frequencies of the motifs were also exploited as classifiers for the selection of the network models [51]. Studying the motifs is also fundamental to understand the stability and robustness of the biological networks to small perturbations [52]. As a matter of fact, Prill *et al.* showed that the stability and robustness of a network is strictly related to the relative abundance of the motifs [52]. This result suggests that the structural organization of a biological network can be highly related to the dynamic properties of the small network motifs. We plan to extend SMGen to incorporate network graphlets and motifs during the generation of the RBMs, to obtain synthetic models with an improved biological significance.

In addition, we will modify the random generation of the kinetic constants to take into account the fact that different biological processes can operate on different time scales. In order to introduce this modification, as a first step the reactions will be grouped into different families based on the biological process that they describe (e.g., protein or mRNA degradation, transcription rates, translation rates). Then, for each family of reactions, a different probability distribution will be used to sample the kinetic constants to reflect the time scale of the described biological process. We are also investigating the possibility of generating models characterized by specific emergent dynamical aspects (e.g., stable oscillatory regimes, state switching, multistability [16]). We derived a set of heuristics to detect these conditions from the time-series, based on the combination of SMGen with GPU-powered stochastic simulators [16]. Still, regardless of the acceleration used, robustly detecting these phenomena is computationally challenging because some parameterizations might lead to very stiff models which, in turn, might require a very long running time. Thus, we are investigating alternative approaches to guide the generation of synthetic models having such features. In particular, we are considering the possibility of providing a set of “functional modules”, i.e, a group of already parameterized reactions implementing specific behaviors. These modules could be coerced into the models generated with SMGen, possibly leading to the emergence of the desired phenomena. Finally, we plan to include an initial check of the parameter values set by the user, based on some heuristics, to verify whether the RBMs can be actually generated as requested. This initial step will allow for avoiding worthless calculations and to suggest useful modifications of parameters to the user.

## Author Contributions

S.G.R., P.C., M.S.N., and A.T. conceived and designed the tool. S.G.R. developed the tool. S.G.R., P.C., and A.T. conceived and designed the analyses. S.G.R. performed the analyses. P.C., D.B., and A.T. wrote the paper. S.G.R., M.S.N., S.S., and L.R. reviewed the paper. D.B., P.C., M.S.N., and A.T. supervised the whole work. All authors have read and agreed to published this version of the manuscript.

## Appendix A. Algorithms

We report here all the algorithms referring to the functions called by Algorithm 1, which represents the workflow of each Worker process.

**Algorithm 2.**
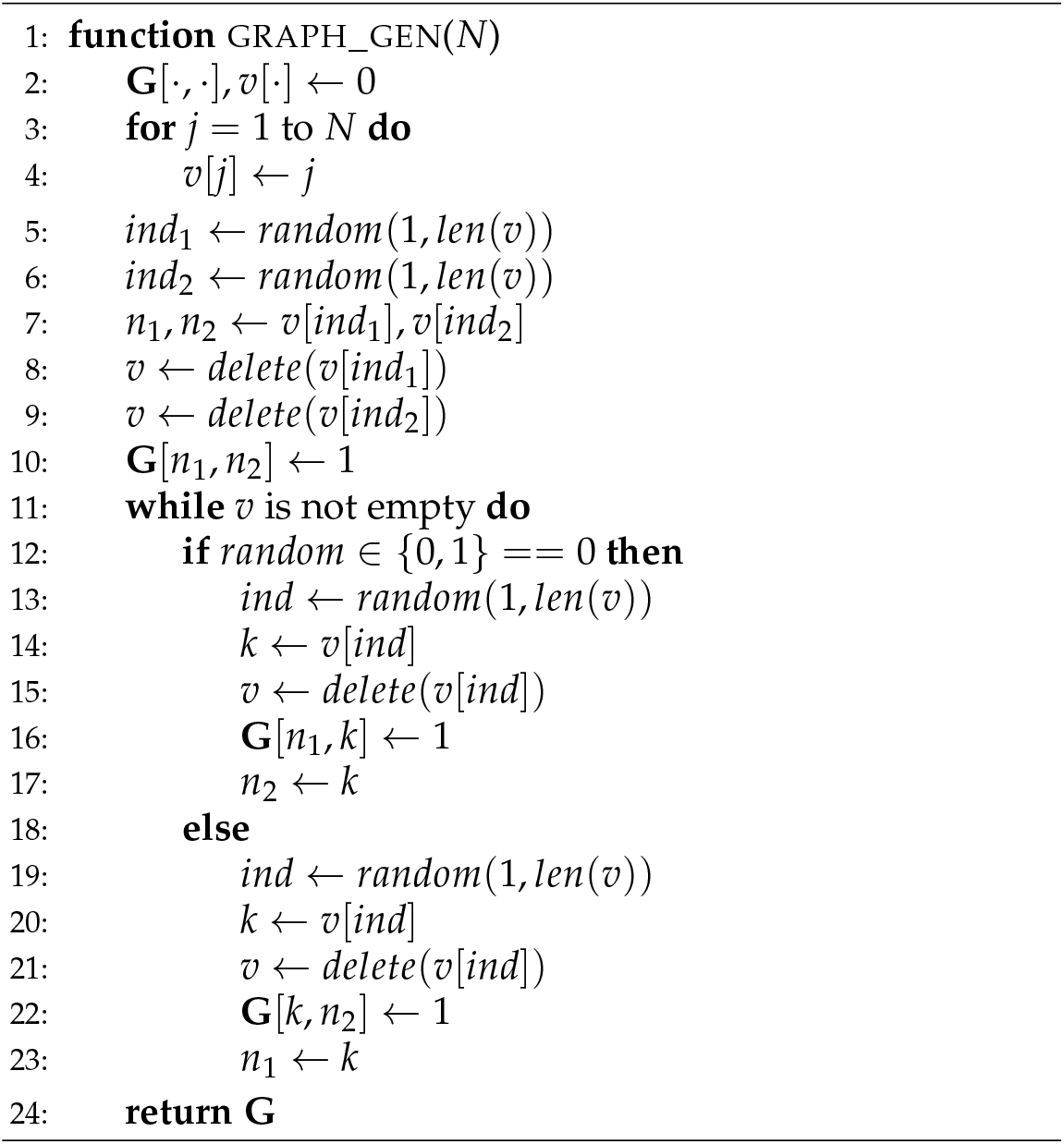
Random initialization of the graph of reactions

**Algorithm 3.**
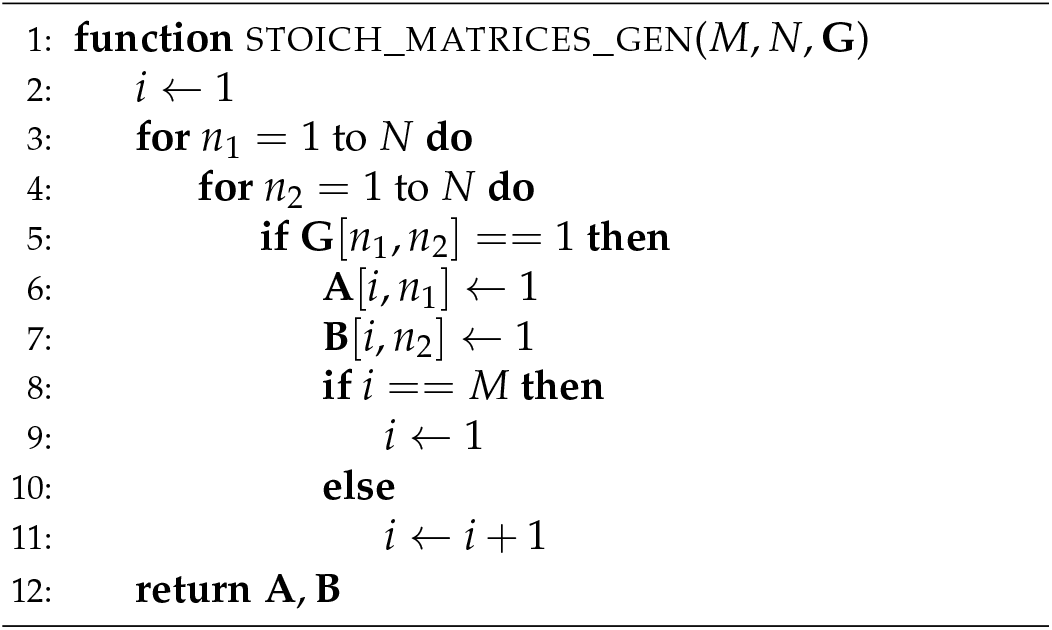
Conversion of the adjacency matrix **G** into the stoichiometric matrices **A** and **B**

**Algorithm 4.**
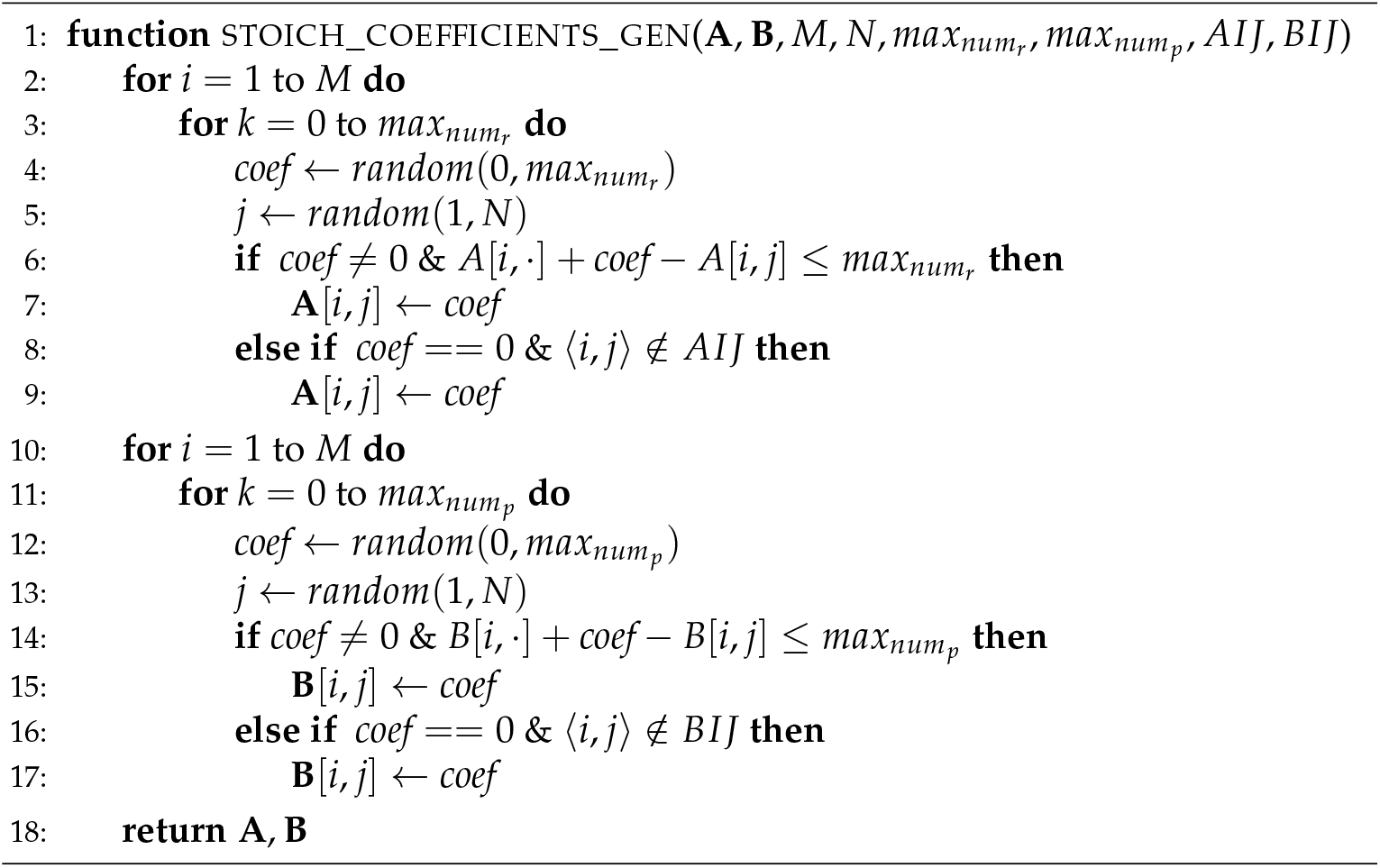
Generation of the random stoichiometric coefficients

**Algorithm 5.**
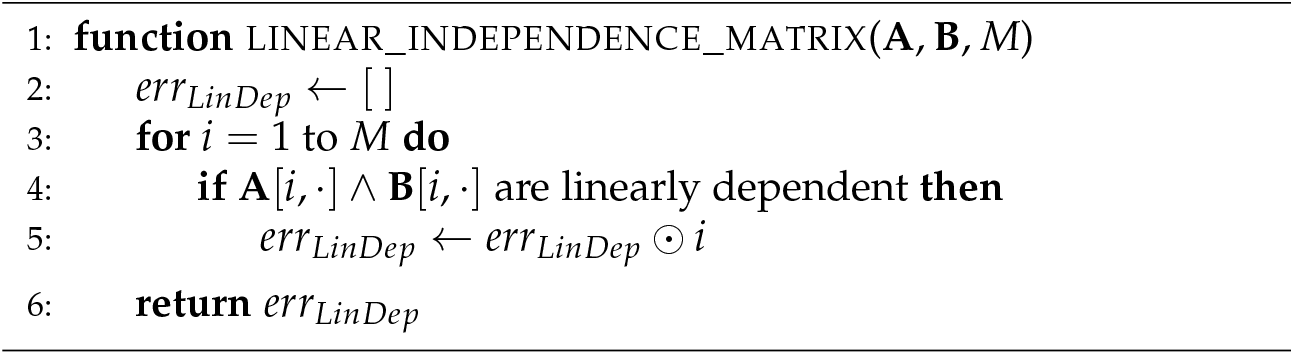
Checking the linear independence between **A**[*i*, ·] and **B**[*i*, ·]

**Algorithm 6.**
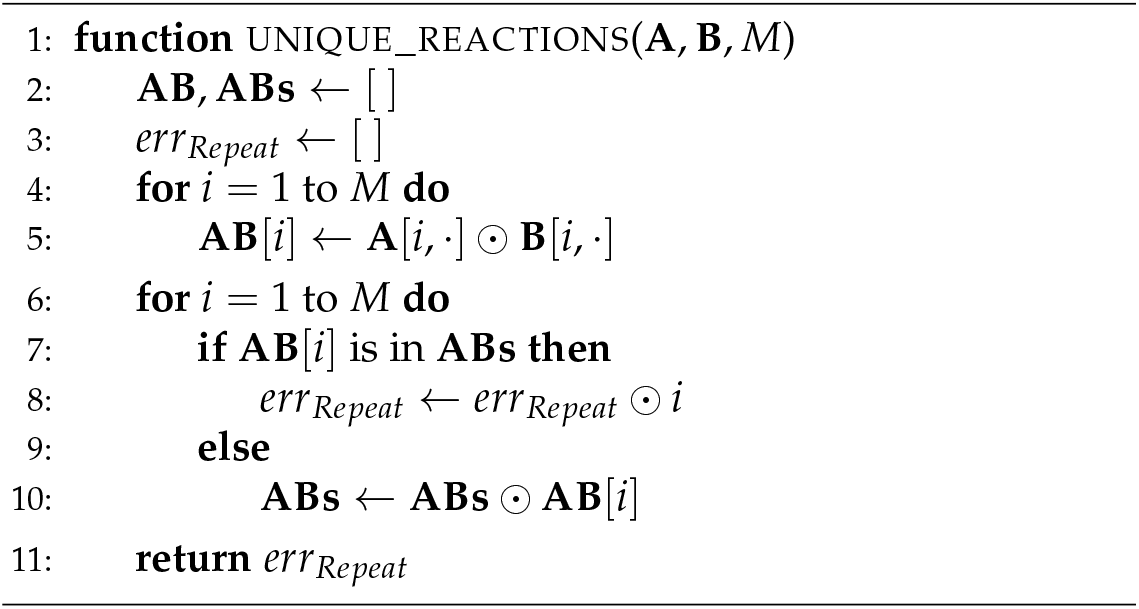
Checking if the generated reactions are unique

**Algorithm 7.**
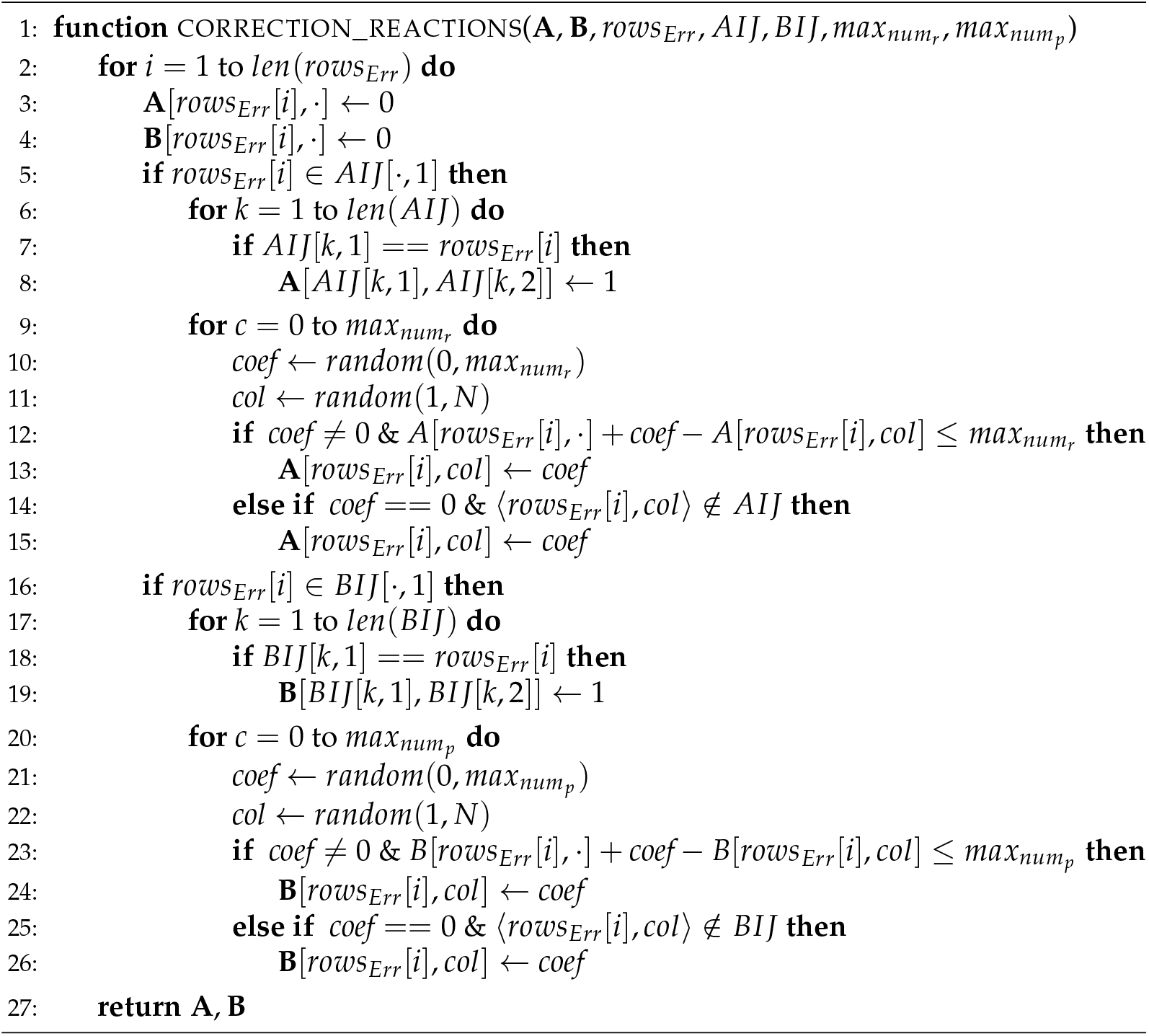
Random correction of the repeated reactions

**Algorithm 8.**
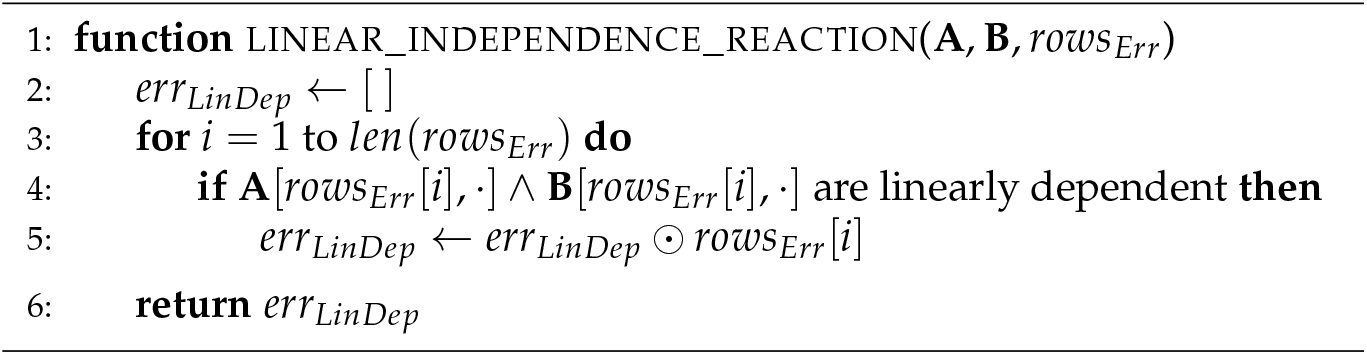
Checking of the linear independence between **A**[*i*, ·] and **B**[*i*, ·]

**Algorithm 9.**
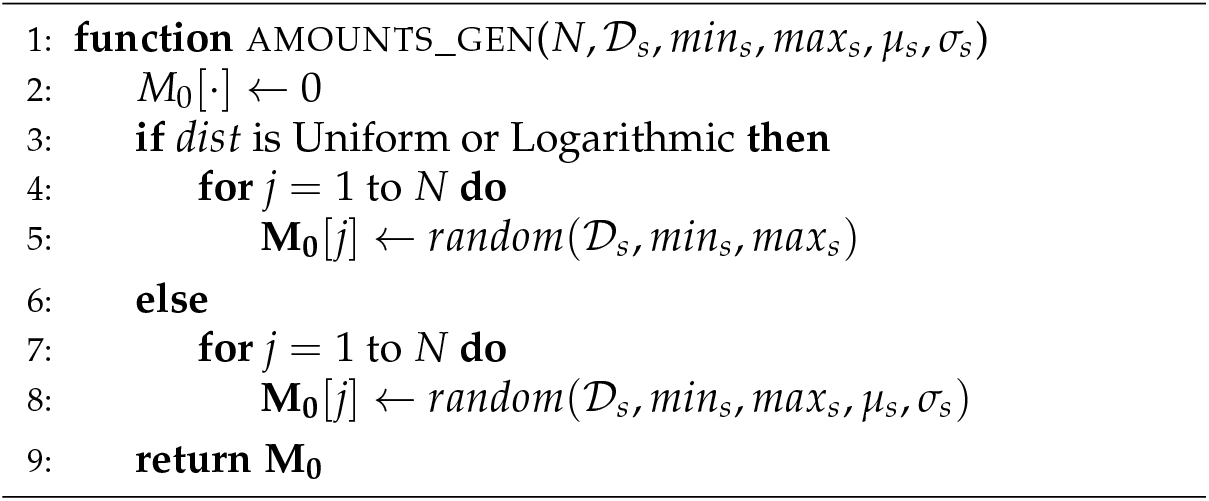
Random initialization of the amounts of the species

**Algorithm 10.**
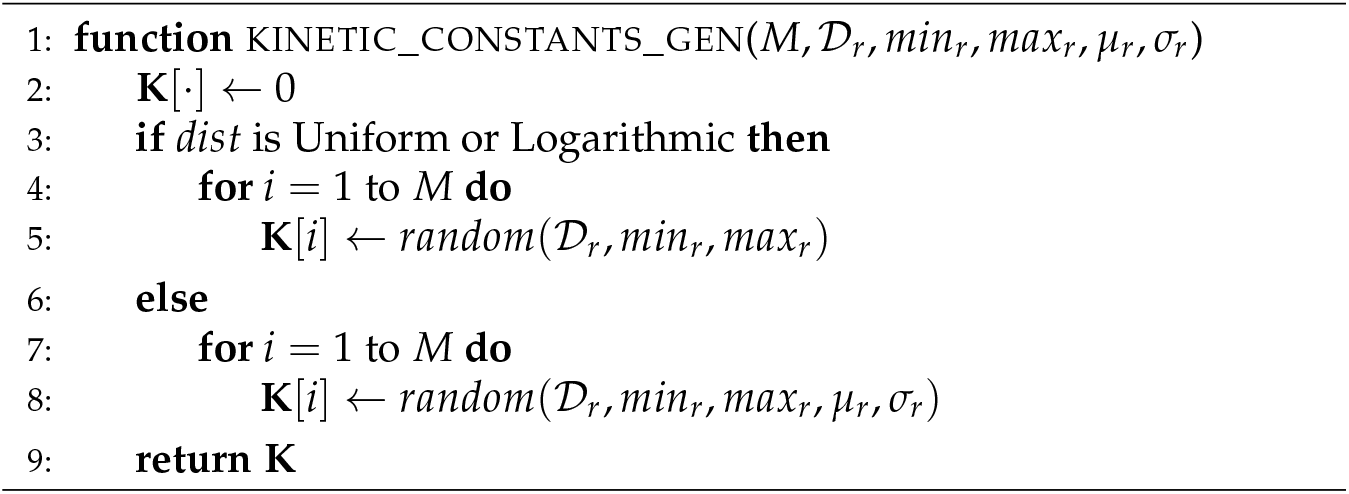
Random generation of kinetic constants of the reactions

**Figure A1.**
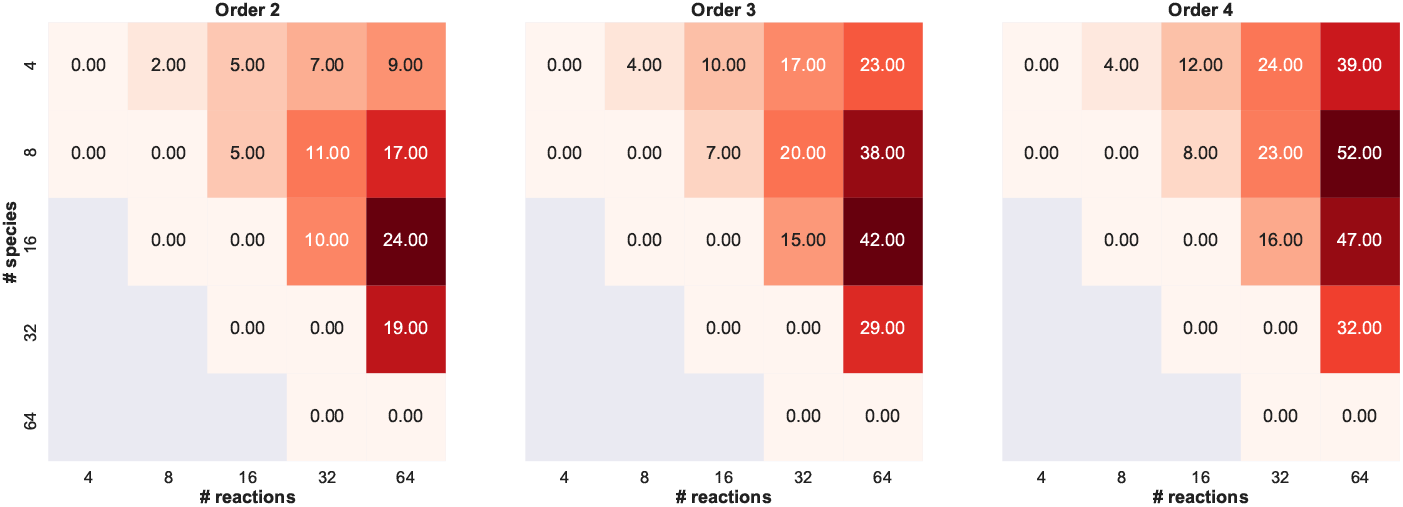
Heatmaps showing the median value of the network deficiency calculated on RBMs generated by varying the number of species and reactions, and the maximum number of reactants and products.

## Appendix B. Analysis of the properties of the RBMs generated with SMGen

We analyzed the properties of the RBMs generated with SMGen by considering the network deficiency [53], the scale-freeness of the generated network [54], and the number of zero complexes, i.e., the null species used to denote degradation reactions or the influx of chemicals.

Network deficiency is an important structural attribute of a reaction network; indeed, this measure gives an indication of the independence of the reaction vectors. Specifically, the network deficiency is a non-negative integer index calculated considering three coefficients: the number distinct complexes n, the number of linkage classes l, and the rank of the network *r.* In this notation, the complexes of a network are the entities that appear at the heads and tails of each reaction (examples of distinct complexes are *A*, 2*A, A* + *B,* etc.), while the linkage classes are the sets of complexes in the various “parts” composing the network, where complexes in a set are linked to each other, directly or indirectly, but they are not linked to any other complex in the network. Finally, the rank of the network is exactly the rank of its set of reaction vectors. To be more precise, the network has a rank *r* if there is a subset containing *r* linearly independent reaction vectors but there is not any subset containing *r* + 1 linearly independent reaction vectors. The network deficiency is equal to *n* – *l* – *r*. It is worth noting that networks having a deficiency equal to zero are the ones where the reaction vectors are as independent as the partition of complexes into linkage classes will allow.

To perform such analysis, as a first step, we generated symmetric and asymmetric RBMs characterized by a number of species *N* and reactions *M* ranging from 4 to 64, by also varying *max_num_r__* and *max_num_p__* into {2,3,4}. Specifically, for each condition (i.e., *N, M, max_num_r__* and *max_num_p__*), 100 RBMs were randomly created to collect statistically sound results. We limited this analysis to the models’ size that can be generated with a single connected component (independently from the maximum number of reactants and products); i.e., with the following constraint:

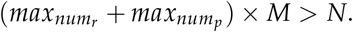

**Figure A2.**
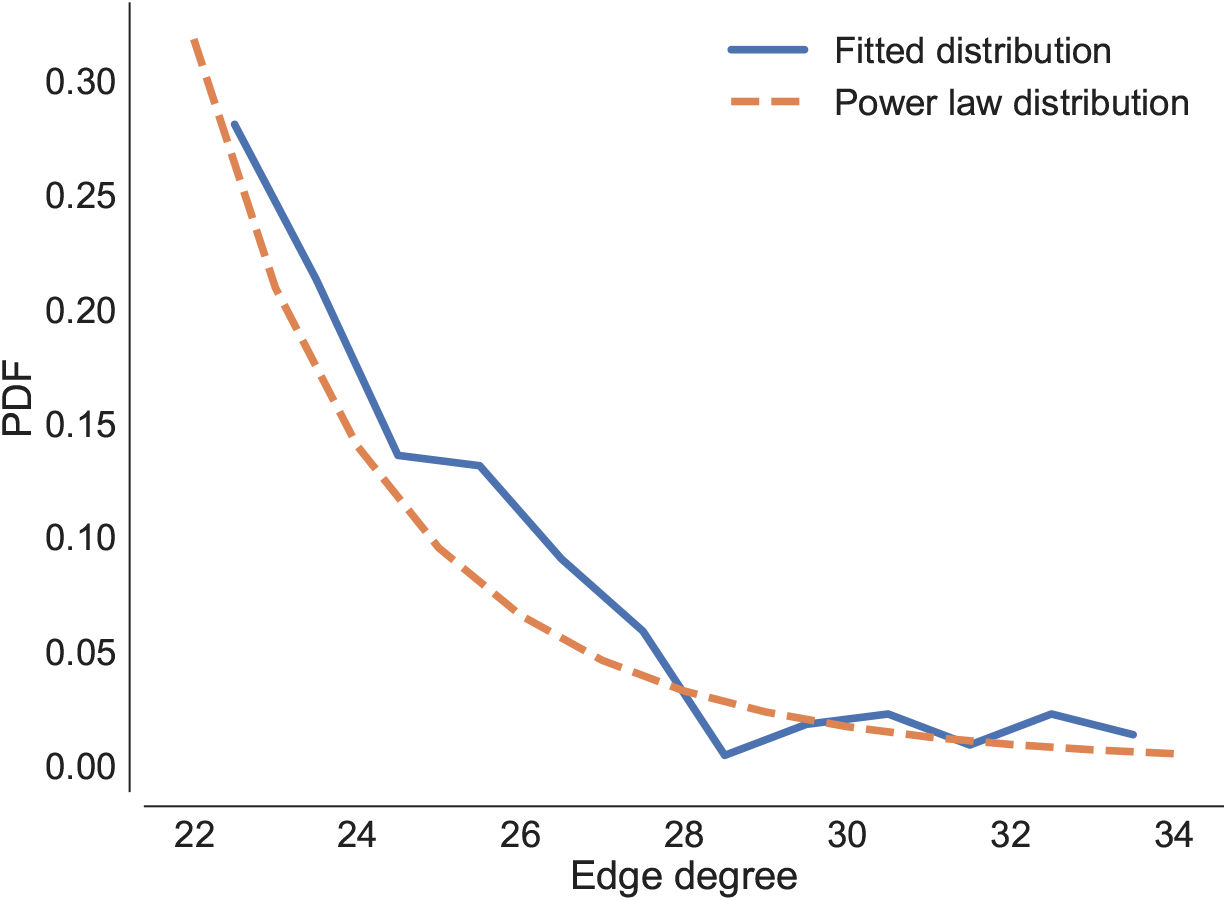
Degree distribution of an interaction network induced by an RBM randomly generated using SMGen. The discrete fitted distribution is shown using linearly spaced bins.

Then, we calculated the network deficiency for each generated RBM; finally, for each condition, we calculated the median of the network deficiency. The heatmaps depicted in Figure A1 clearly show that the network deficiency increases when *M > N.* Moreover, the maximum number of reactants and products affects the network deficiency: indeed, the higher the maximum number of reactants and products, the higher the network deficiency. The achieved results are coherent with the definition of network deficiency itself. As a matter of fact, the number of possible distinct complexes and the number of possible distinct linkage classes increase along with *N* and the maximum number of reactants and products, while the number of linearly independent reaction vectors is bounded by min(*M, N*). Finally, the results obtained from this analysis present values congruent with those of real biochemical models, such as the Brussellator [38,43]) (5 species and 4 reactions) with deficiency = 0, the prokaryotes gene expression model [55] (5 species and 8 reactions) with deficiency = 1, the heat shock response in eukaryotes model (10 species and 17 reactions) with deficiency = 5, the Ras/cAMP/PKA model [22] (33 species and 39 reactions) with deficiency = 7, and the human intracellular metabolic pathway in red blood cells [56] (114 species and 226 reactions) with deficiency = 13.

Biological networks, in particular metabolic networks [57], often show scale-freeness properties [54]. In order to prove that SMGen produces networks that are characterized by this property, we converted the bipartite graph defined by the stoichiometric matrices to a common interaction graph, where nodes represent the chemical species and edges represent reactions involving the nodes as reactants or products. Then, we analyzed the structural properties of the generated network and, in particular, we investigated whether the degree distribution follows a power law distribution, which is characteristic of scale-free graphs. Since the scale-free properties emerge only for large scale networks, we generated and analyzed the interaction network induced by an RBM with by 10,000 chemicals species involved in 10,000 reactions. According to our results, the degree distribution seems to fit well with a power law (see Figure A2), confirming that SMGen can generate scale-free networks. It is worth noting that curated models of this size are seldom published, because both the initial amounts of the chemical species and the kinetic reaction rates would be difficult to collect, and in any case they would represent a challenge for biochemical simulation.

**Figure A3.**
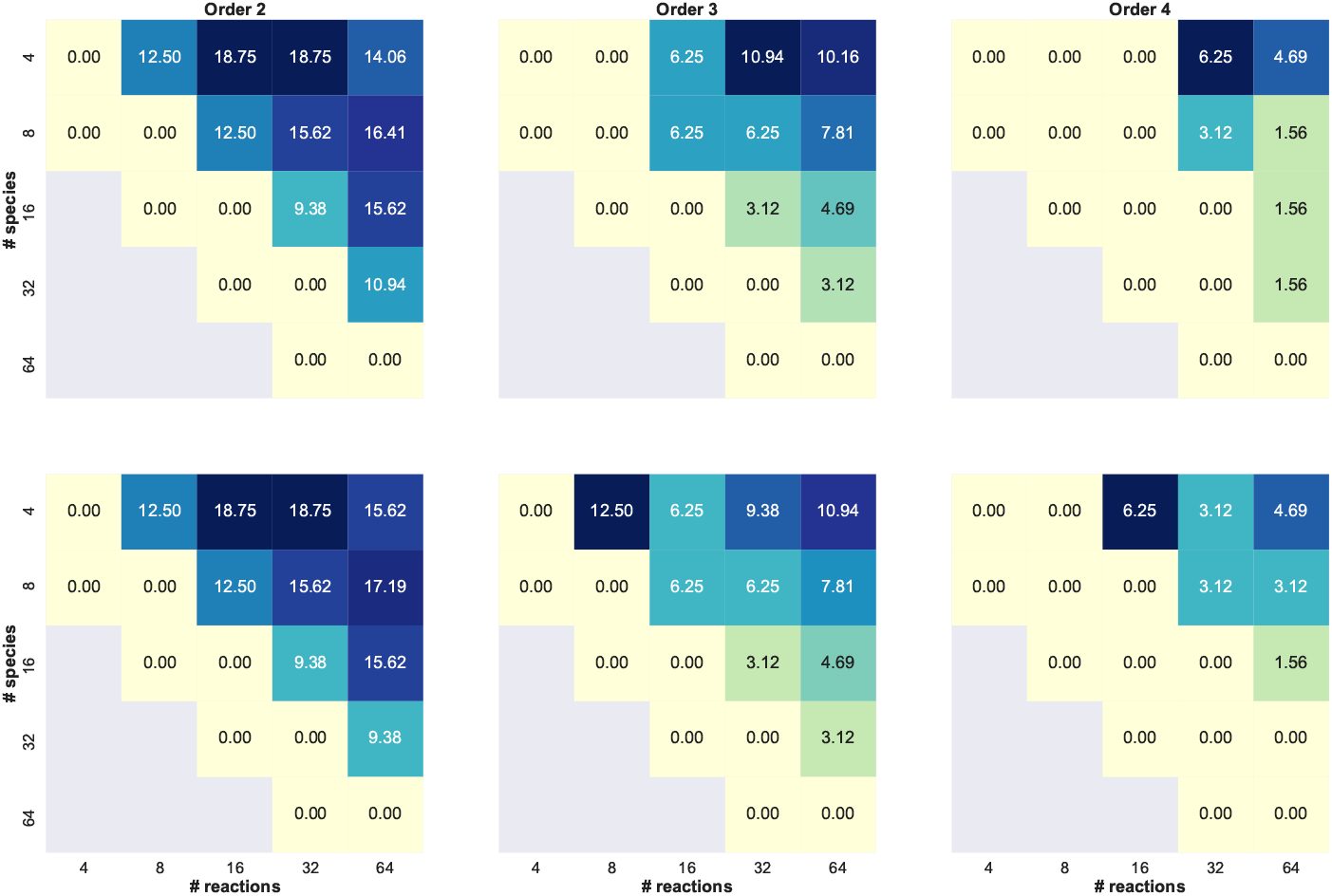
Heatmaps showing the median percentage of reactions where the zero complex appears as reactant (top) or product (bottom), calculated on RBMs generated by varying the number of species and reactions, and the maximum number of reactants and products

Finally, in order to evaluate the abundance of the zero complex appearing as reactant or product in a reaction, we used the same RBMs generated to calculate the network deficiency. As a first step, for each RBM, we calculated the percentage of reactions where the zero complex appears either as a reactant or as a product. The heatmaps reported in Figure A3 clearly show that this value increases when *M* >> *N*. It is worth noting that the higher the maximum number of reactants and products for each reaction, the lower the percentage of reactions involving the zero complex. Overall, we observe that at most 2 out of 10 species are involved in one of such reactions. We investigated the abundance of the zero complex in real models. In the prokaryotes gene expression model [55], the zero complex appears in 25% of the reactions, while it is present in 5.88% of the reactions composing the heat shock response model [58]. Thus, the values calculated on synthetic RBMs generated by SMGen are in agreement with those obtained by analyzing models representing real biochemical networks.

